# Methicillin-Resistant Staphylococcus Aureus CC22-MRSA-IV as an Agent of Dairy Cow Intramammary Infections

**DOI:** 10.1101/326553

**Authors:** Giada Magro, Marta Rebolini, Daniele Berettac, Renata Piccinini

## Abstract

Methicillin-resistant *S. aureus* (MRSA) lineages have become major responsible of healthcare-and community-associated infections in human population. Bovine MRSA are sporadically detected in the dairy herd, but its presence might enhance the risk of zoonosis. Some lineages are able to lose the specific host tropism, being easily transmitted from animals to humans and vice-versa. The present study aims at clarifying the epidemiology of MRSA intramammary infections in a closed dairy herd, which was running a mastitis control program since years. Quarter milk samples were collected from all lactating cows once a week for 9 weeks and bacteriologically tested. At the end of the follow-up period, also a self taken nasal swab of the milker was analysed. Three cows (12.5%) were MRSA positive, a four showed a transient infection and an MRSA was isolated also from the milker. Somatic cell counts of the infected quarters fluctuated from 1,000 to 1,800,000 cells/mL. All isolates were genotyped using DNA microarrays and identified as the epidemic UK-EMRSA-15 grouping in CC22. All strains carried the genes for β-lactam and macrolide resistance. The milker isolate differed from cow isolates mainly for the absence of the untruncated β-haemolysin and the presence of the immune evasion cluster. The milker had been volunteering in a nursing home since months, thus playing the role of MRSA vector into the herd. Our results showed the adaptive capacity of such MRSA to the bovine host. Therefore, we suggest that CC22-MRSA should be regarded as a potential cause of humanosis in dairy cattle herds.

## IMPORTANCE

Animals are the major source of new pathogens affecting human populations. However, the potential for pathogenic bacteria originally isolated in humans, to switch hosts and adapt to mammals is not to underestimate. Here, we report the emergence and spread of subclinical intramammary infections caused by a methicillin-resistant *Staphylococcus aureus* of human origin, in a closed dairy herd. The strain, responsible for epidemics in England and other Countries, was isolated also from the milker’s nose, suggesting a host-adaptive evolution inside the herd. Our findings demonstrate that the human worker can act as a reservoir for contagious *Staphylococcus aureus* clones with potential for herd spread, highlighting the need of considering also the risk of humanosis in *Staphylococcus aureus* mastitis control programs.

## INTRODUCTION

*Staphylooccus aureus (S. aureus)* is widely known as the major cause of contagious bovine mastitis and an important pathogen in different livestock species^1^. The treatment with β-lactam antibiotics resulted in a selective pressure for resistance, and the acquisition of the mobile staphylococcal cassette chromosome *(SCCmec)*, carrying the *mecA* or *mecC* gene, allows the bacteria to continue the cell wall biosynthesis, nullifying the antibiotic action. Methicillin-resistant *S. aureus* (MRSA) lineages are the result of this successful evolution, becoming a major responsible of healthcare- and community-associated infections on a global scale^2^. In contrast with the human-associated lineages, bovine MRSA are sporadically detected in the dairy herds, being mostly associated with low prevalence of subclinical mastitis. Despite that, the persistence of MRSA clones in dairy herds might enhance the risk of zoonosis^3^. From the first bovine MRSA detected about 50 years ago^4^, understanding the risk of *S. aureus* cross-species transmission is still an interesting scientific field of research. The phylogenetic studies on MRSA demonstrated that bovine strains belong to a limited group of clonal complexes (CC)^5,6^. Human lineages of MRSA, such as CC5, CC8, CC22, CC30 and CC45 are rarely found in animals, suggesting host range barriers^7,8^. On the animal side, the most common livestock isolates belong to a small number of animal-associate clones: in particular bovine mastitis isolates group in few CCs, including CC1, CC8, CC97, CC126, CC130, CC133, CC398 and CC705^1^. Some of these have been demonstrating their ability to shift from animal to human hosts. This is the case of CC398 MRSA: firstly isolated in pig, poultry and ruminant farms, it is now displaying a zoonotic potential, contributing to the MRSA widespread diffusion in the human healthcare system^9^. By contrast, CC8 originated in humans and emerged in the cow after ancient or recent host jumps^10^. The new bovine-adapted genotype loses the ability to colonize humans, lacking of a human-related mobile genetic element^11^. Therefore, if some *S. aureus* clones can lose the specific host tropism and be easily transmitted from animals to humans and vice-versa, we need to expand the concept of zoonosis including also humanosis. This study aims at clarifying the epidemiological origin of a new MRSA intramammary infection in a closed dairy cow herd, which was running a mastitis control program since years.

## RESULTS

The results of bacteriological analysis of quarter milk samples collected at the first sampling showed that 3 out 24 lactating cows (12.5%) had 2 up to 3 quarters infected by *S. aureus.* PCR assay confirmed the identification of the isolates as *S. aureus.* During the follow-up period, SCC of the infected quarters fluctuated from extremely low values (1,000 cells/mL) to values exceeding one million cells/mL. At the third sampling, another animal tested positive in one quarter, but cured spontaneously within 3 weeks and remained negative in the following two months (the quarter was tested repeatedly until the end of August). The cow showed always very low SCC, never exceeding 7,000 cells/mL. Two infected animals were culled before the end of the study, i.e. after the 7^th^ or 8^th^ sampling respectively. Somatic cell count values and *S. aureus* shedding by the infected quarters of the 4 cows are presented in FIG 1.

**FIG 1.**
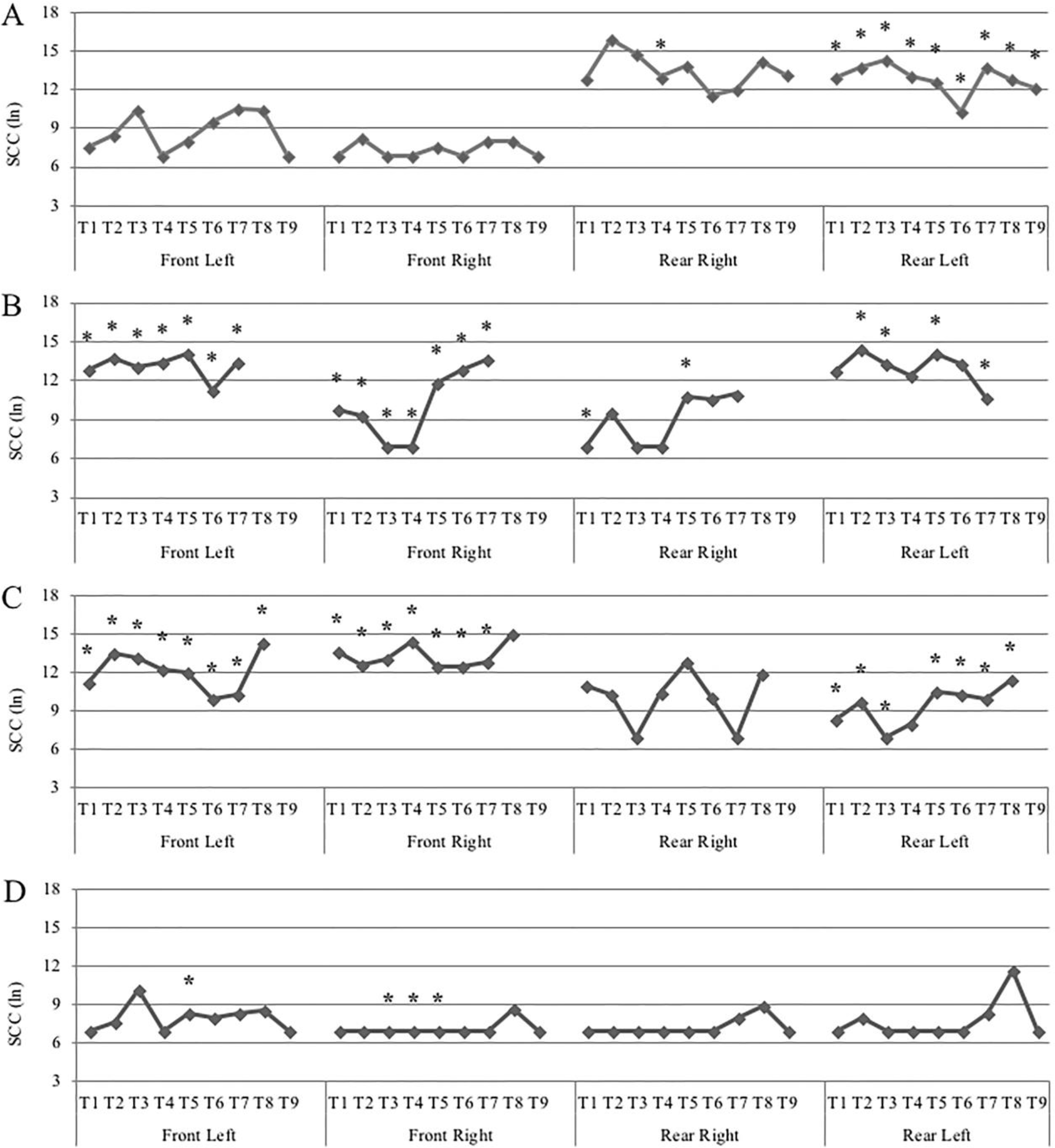
Quarter milk Somatic Cell Counts and MRSA shedding by infected cows during the study. The capital letters A-D indicate the four cows. The symbol * represents the recovery of *S. aureus* in the milk.

*S. aureus* was recovered also from the milker’s nasal swab.

The disk diffusion test showed the same pattern of antibiotic resistance for all *S. aureus* isolates: they were susceptible to macrolides and rifaximin, but resistant to penicillin, ampicillin, amoxicillin/clavulanate, oxacillin, 1^st^, 3^rd^ and 4^th^ generation cephalosporins, kanamycin and quinolones. Therefore, the isolates were classified as MRSA.

Microarray genotyping evidenced the *mecA* gene in all the 5 isolates, including the human one. They were identified as epidemic MRSA-15 (also known as UK-EMRSA-15 or Barnim EMRSA) and grouped in CC22. The microarray results showed minor differences among the isolates, as reported in TABLE 1. All cow isolates carried the γ-haemolysin genes *hlgA* and *hlgB*, only the strain isolated from the last infected cow carried *hlgC.* All isolates were Panton-Valentine leucocidin (PVL) negative, but positive for the enterotoxin genes *seg, sei, sem, sen, seo* and *seu* an allelic variant of von Willebrand factor (vvb-RF122). They harboured also the protease genes encoding aureolysin or staphopain A, B (data not shown). Human and cow isolates differed basically for the absence of the untruncated β-haemolysin and the presence of *sak*, *chp* and *scn* uniquely in the milker *S. aureus.* The demonstration of the genes for β-lactams resistance in all isolates explained the phenotypic resistance observed. Conversely, *ermC*, one of the genes encoding macrolide resistance, did not express resistance to tylosin or sipiramycin in the susceptibility test.

**Table 1.**
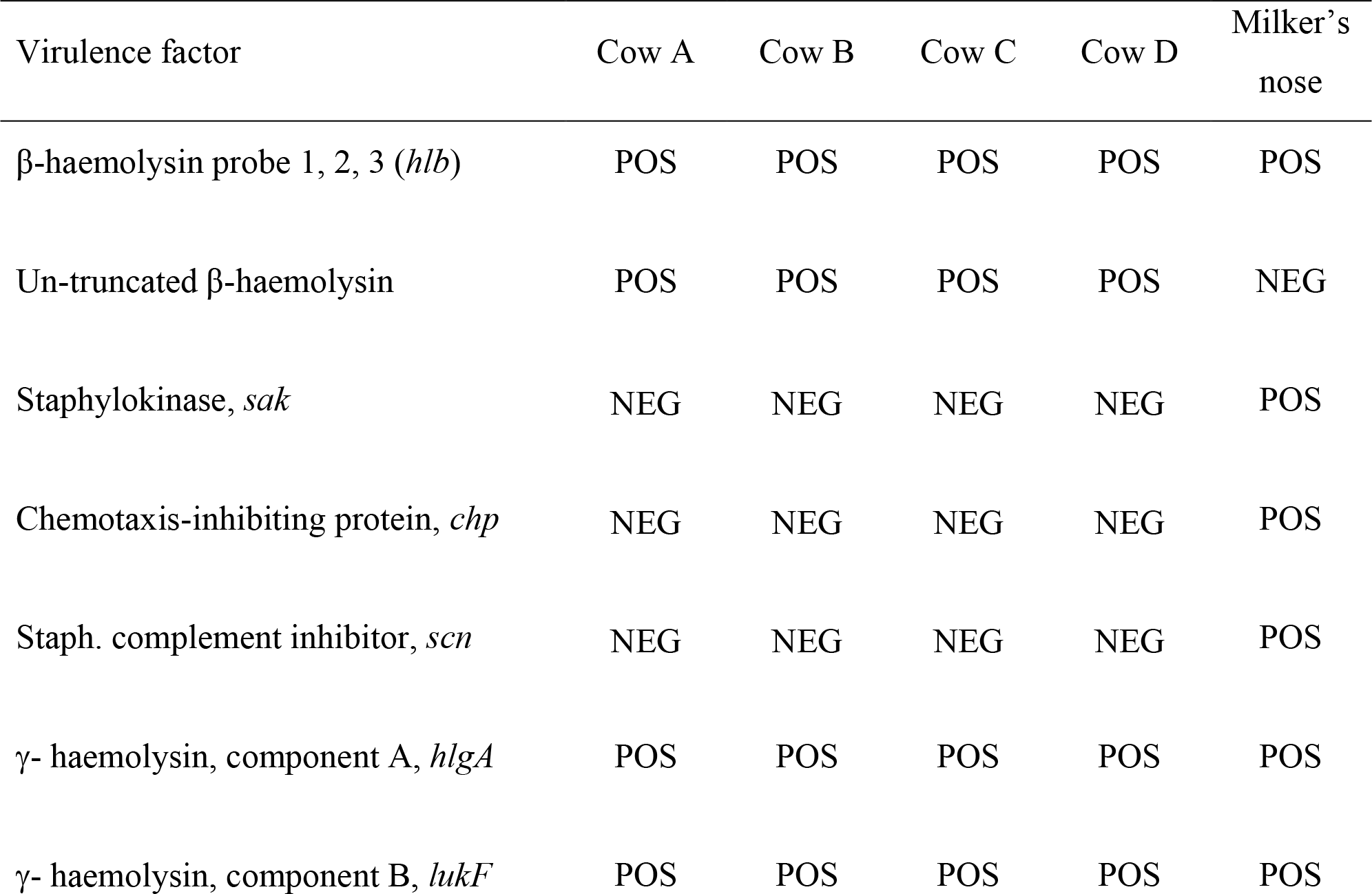
Main results of microarray analysis showing the differences among the MRSA
strains isolated during the study.

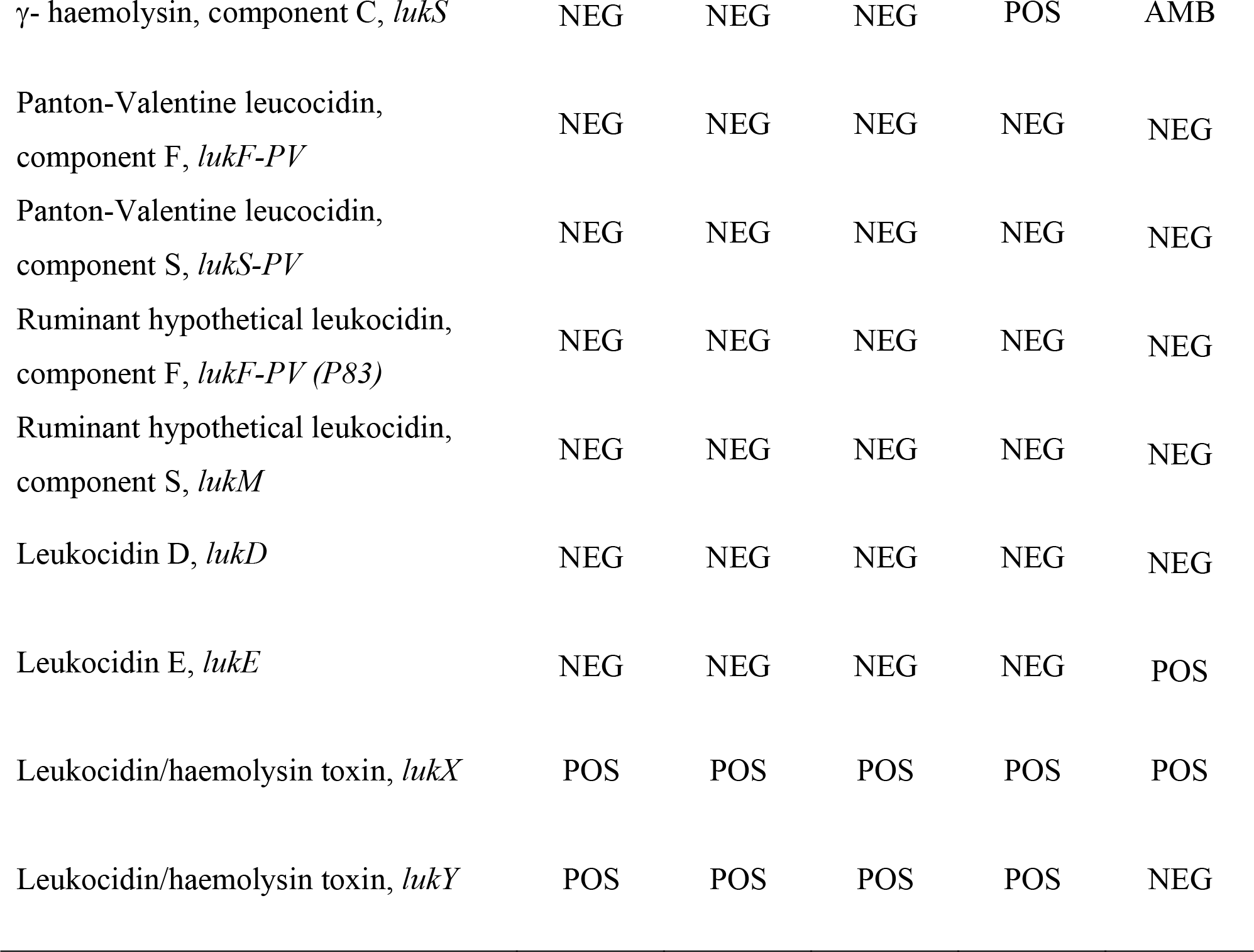

## DISCUSSION

Methicillin-resistant *S. aureus* (MRSA) strains are the major cause of healthcare- and community-associated infections on a global scale^2^. Different lineages, termed as livestock-associated MRSA (LA-MRSA) are implicated in farm animal infections. The possible transmission of human lineages to companion animals, through their owners or caretakers is widely demonstrated^12,13,14^, therefore the infection is regarded as a humanosis. In dairy cattle, MRSA is usually considered as a marginal problem in terms of herd contagiousness but at the same time, a possible reservoir of new human infection^3^. Conversely, the concept of humanosis is still poorly considered. The reason behind this underestimation is probably due to the difficult demonstration of the epidemiological chain leading to the infection in the intensive dairy herd, what makes the distinction between zoonosis and humanosis a complicated problem. In the last decades, several studies focused on the possible transmission of LA-MRSA to human population, demonstrating the zoonotic role of some lineages in pig, cattle, and poultry farm workers^15,16^. CC398 is the most important group and the possible colonization of cattle farm personnel has been considered as a potential MRSA vector into different compartment of the farm^17^ or into hospital^18^. The results of the present study led us to consider the subclinical intramammary infections of the dairy cows not as a zoonosis, but a humanosis, since all *S. aureus* isolates from quarter milk and the isolate from the milker’s nose belonged to the same clonal lineage, i.e. the epidemic UK-EMRSA-15. It should be highlighted that the milker volunteered since months in a nursing home. Such lineage is largely diffused in pets: dogs and cats acquire the infection by their owners or veterinarian^19^. The genome comparison of CC22-MRSA isolated from humans and pets demonstrated a few differences, mostly in the carriage of mobile genetic elements (MGEs) rather than in core genes^14^. Indeed, the lineage is characterized by a flexible MGEs profile, associated with a quick ability of MGEs loss and acquisition, which might explain its success in dissemination and persistence in different hosts^20^. In our study, the microarrays results showed some differences in the occurrence of the immune evasion cluster (IEC): the β-haemolysin converting prophage carrying human-specific host immune evasion genes *(sak-scn-chp)* was present only in the human isolate, suggesting a quick adaptation of the lineage to the bovine host. This finding is similar to the case of CC8 human-to-animal jump^10,11^. Analogously to CC8, the loss of the prophage might help the establishment of infection in the dairy cow. A further result strengthening our hypothesis is the presence of the untruncated β-haemolysin uniquely in the bovine MRSA isolates, probably because the gene is necessary in ungulates for the different structure of erythrocyte membranes. The outbreak and dissemination of CC22-MRSA infection in the herd before our monitoring support the hypothesis that the adaptation of the lineage to this new host should not be underestimated. The cow D isolate differed from the other bovine isolates for the carriage of the *hlgC/lukS* gene, which in turn gave an ambiguous result in the human isolate. We would like to highlight this result, because the cow was the only one affected by a transient intramammary infection. We could speculate that the pathogenicity island carrying γ-haemolysin might have been lost in the adaptation to the bovine host. All the isolates harboured the allelic variant of the Von Willebrand binding protein gene (*vvb-RF122)*, which is considered one of the mechanisms associated to *S. aureus* pathogenicity in the cow and a specific marker of host adaptation^21^. At the light of these results, we strongly suggest that CC22-MRSA be regarded as a potential cause of humanosis in dairy cattle herds.

## Conclusions

The present study provides evidence for the importance and impact of the UK-EMRSA-15 as a cause of mastitis in the dairy cow, demonstrating the adaptive capacity of the lineage to the bovine host. The transmission of MRSA between different hosts revoke the concept of “One Health”: the true scale of the problem is still unknown, and further studies addressing both animals and farm personnel are required, in order to monitor the possible emergence of new lineages among the dairy cattle. In order to minimize the risk of *S. aureus* spread within-herd and in the community, the herd biosecurity measurements should be implemented.

## MATERIAL AND METHODS

### Herd history

The study was performed in a small farm located in Lombardy region. The herd is housed in freestall with cubicle barns and milling parlour. A contagious mastitis control program has been running since years, because raw milk is sold directly at the farm. One year and half before our study, the routine bacteriological analysis of bulk tank milk had evidenced the presence of *S. aureus*, with a value of 40 CFU/mL. Quarter milk samples were collected from all the cows and the new infected ones were milked after the healthy animals, but not physically segregated. After 6 months, *S. aureus* count had increased to 140 CFU/mL. Therefore, the owner decided to cull part of the infected animals, so that 6 months before the beginning of the present study the bulk milk concentration of *S. aureus* had decreased to 73 CFU/mL. New cows were not introduced into the herd, therefore the total number of lactating animals was 24.

### Sampling and bacteriological analysis

Quarter milk samples of all the lactating animals were aseptically collected once a week for 9 weeks (T1 to T9) during milking in the months of April to June, and immediately delivered to the laboratory. Bacteriological analysis was performed as previously indicated ^22^ and somatic cells (SCC) were counted using a Bentley Somacount 150 (Bentley, USA).

At the end of the follow-up period, we also analysed a self-taken nasal swab of the milker. The isolates were presumptively identified as *S. aureus* according to the following scheme: Gram-positive cocci, haemolytic on blood agar, catalase positive, and coagulase positive in 4–24 h.

The antibiotic resistance to the drugs mostly used in mastitis therapy (penicillin, ampicillin, amoxicillin/clavulanate, oxacillin, 1^st^, 3^rd^ and 4^th^ generation cephalosporins, tylosin, kanamycin, rifaximin, quinolones, thiamphenicol, trimethoprim/sulfamethoxazole) was tested by disk-diffusion.

### Molecular analysis

The DNA of coagulase-positive strains was extracted using DNeasy kit (QIAgen, Hilden, Germany), with the addition of lysostaphin (5 mg/mL; Sigma-Aldrich, St. Luis, MO, USA) for bacterial lysis. Amount and quality of DNA samples were measured on a NanoDrop ND-1000 spectrophotometer (Nano-Drop Technologies, Wilmington, DE, USA).They were confirmed as *S. aureus* by a duplex real-time PCR assay^23^.

Genotyping was performed by DNA microarrays using Alere StaphyType DNA microarray (Alere Technologies Gmbh, Jena, Germany). The microarray covers approximately 170 distinct genes and their allelic variants for a total of 330 target sequences including accessory gene regulator alleles, genes coding for virulence factors and for microbial surface components recognizing adhesive matrix molecules (MSCRAMMs), capsule type-specific genes, and numerous antimicrobial resistance genes^24^. Probes for the methicillin-resistance genes *mecA* and *mecC* are also included. The overall pattern was analyzed automatically for the presence or absence of specific genes and compared to a database of strain profiles allowing the assignment to Clonal Complexes (CC). The genotyping service was performed at Alere Technologies (Jena, Germany).

## ACKNOWLEDGMENTS

We thank L. Zanini, milk specialist of the Breeder Association of Lombardy, for supplying
the bacteriological results of previous analysis of bulk milk.

